# Avian thermoregulation in the heat: is evaporative cooling more economical in nocturnal birds?

**DOI:** 10.1101/282640

**Authors:** Ryan S. O’Connor, Ben Smit, William A. Talbot, Alexander R. Gerson, R. Mark Brigham, Blair O. Wolf, Andrew E. McKechnie

**Author notes:** **Summary statement:** Caprimulgids and Australian owlet-nightjars displayed allometrically lower water losses compared to diurnal birds, whereas owls exhibited water losses comparable to similarly sized diurnal birds.

## Abstract

Evaporative cooling is a prerequisite for avian occupancy of hot, arid environments, and is the only avenue of heat dissipation when air temperatures (T_a_) exceed body temperature (T_b_). Whereas diurnal birds can potentially rehydrate throughout the day, nocturnal species typically forgo drinking between sunrise and sunset. We hypothesized that nocturnal birds have evolved reduced rates of evaporative water loss (EWL) and more economical evaporative cooling mechanisms than those of diurnal species that permit them to tolerate extended periods of intense heat without becoming lethally dehydrated. We used phylogenetically-informed regressions to compare EWL and evaporative cooling efficiency (ratio of evaporative heat loss [EHL] and metabolic heat production [MHP]; EHL/MHP) among nocturnal and diurnal birds at high T_a_. We analyzed variation in three response variables: 1) slope of EWL at T_a_ between 40 and 46°C, 2) EWL at T_a_ = 46°C, and 3) EHL/MHP at T_a_ = 46°C. Nocturnality emerged as a weak, negative predictor, with nocturnal species having slightly shallower slopes and reduced EWL compared to diurnal species of similar mass. In contrast, nocturnal activity was positively correlated with EHL/MHP, indicating a greater capacity for evaporative cooling in nocturnal birds. However, our analysis also revealed conspicuous differences among nocturnal taxa. Caprimulgids and Australian-owlet nightjars had shallower slopes and reduced EWL compared to similarly-sized diurnal species, whereas owls had EWL rates comparable to diurnal species. Consequently, our results did not unequivocally demonstrate more economical cooling among nocturnal birds. Owls predominately select refugia with cooler microclimates, but the more frequent and intense heat waves forecast for the 21^st^ century may increase microclimate temperatures and the necessity for active heat dissipation, potentially increasing owls’ vulnerability to dehydration and hyperthermia.

## Introduction

Maintaining heat balance can present a significant physiological challenge for animals, particularly when inhabiting very hot environments (Porter and Gates, 1969). When environmental temperature exceeds body temperature (T_b_), animals experience a net heat gain and evaporative water loss (EWL) becomes the only mechanism whereby they can dissipate internal heat loads (Dawson, 1982; Tattersall et al., 2012). The quantity of water lost, relative to body mass (M_b_), can become substantial at high air temperatures (T_a_) in small birds (McKechnie and Wolf, 2010). For example, EWL can exceed 5% of M_b_ hr^-1^ in Verdins (*Auriparus flaviceps*) experiencing 50°C temperature (Wolf and Walsberg, 1996). Therefore, during extreme heat waves, the thermoregulatory requirements necessary to avoid lethal hyperthermia could exceed physiological limits of birds and other animals, an occurrence that sometimes leads to large-scale mortality events (Welbergen et al., 2008; McKechnie et al., 2012; Fey et al., 2015).

Like many physiological traits, a large proportion of the variation in avian EWL can be attributed to scaling with M_b_ (Bartholomew and Dawson, 1953; Crawford and Lasiewski, 1968). However, substantial mass-independent variation remains among similarly-sized species (Lasiewski and Seymour, 1972, Smit et al., 2017) and understanding the sources and significance of this variation is a central focus in the fields of ecological and evolutionary physiology. Climate has frequently been invoked as an ultimate driver of adaptive variation in avian EWL (Williams, 1996; Tieleman et al., 2002). For instance, measurements of EWL at moderate T_a_ (e.g., 25°C) suggest that arid-zone species have evolved reduced rates of water loss compared to their mesic counterparts (Williams and Tieleman, 2005). Although fewer data exist for hot conditions, rates of water loss at T_a_ > T_b_ exhibits similar trends, at least intraspecifically, with EWL rates lower in arid-zone populations compared to those inhabiting more mesic habitats (Trost, 1972; Noakes et al., 2016; O’Connor et al., 2017). Climate and M_b_ scaling may not, however, be the only ecological factors influencing variation in EWL (e.g., diet; Bartholomew and Cade, 1963).

One additional factor that could conceivably exert a strong influence on evaporative cooling economy is the timing of activity (i.e., nocturnality *versus* diurnality). Diurnal birds can potentially gain water through drinking and/or foraging during the course of the day, thereby offsetting water losses via evaporation (Willoughby and Cade, 1967; Fisher et al., 1972). In contrast, nocturnal birds often experience zero water gain aside from metabolic water for as long as 12 - 14 hours between sunrise and sunset (Brigham, 1991). Consequently, if daytime environmental temperature exceeds T_b_, roosting nocturnal birds are subjected to extended periods of elevated requirements for evaporative cooling (Grant, 1982). Birds from the family Caprimulgidae (nightjars and nighthawks) have frequently been reported roosting/nesting in unshaded sites where environmental temperature can approach 58°C (Cowles and Dawson, 1951; Weller, 1958; Grant, 1982; Ingels et al., 1984). Even owls, which typically select thermally buffered refugia, can experience T_a_ approaching or exceeding T_b_ in hot habitats (Ligon, 1968). For example, Pearl-spotted Owlets (*Glaucidium perlatum*) and African Scops Owls (*Otus senegalensis*) occur in the Kalahari Desert of southern Africa, and Western Screech Owls (*Megascops kennicottii*), Elf Owls (*Micrathene whitneyi*) and Ferruginous Pygmy-Owls (*Glaucidium brasilianum*) occur year-round in the Sonoran Desert, where midsummer T_a_ maxima can approach 50°C (Wolf, 2000). Thermal trade-offs involved with site selection may also be important; Rohner et al. (2000) observed Great Horned Owls (*Bubo virginianus*) shifting roost sites seasonally to more open, thermally exposed areas during summer, a behavior attributed to black fly avoidance. Given these behavioral constraints, we predict that nocturnal species in hot environments have experienced strong selection for efficient evaporative cooling and reduced EWL to minimize dehydration risk during the day.

A parameter often used to quantify thermoregulatory capacity at T_a_ > T_b_ is evaporative cooling efficiency, calculated as the ratio of evaporative heat loss and metabolic heat production (EHL/MHP; Lasiewski et al., 1966). An EHL/MHP value greater than one signifies that an individual can dissipate all its metabolic heat through evaporation, with higher EHL/MHP ratios indicating more efficient evaporative cooling. When evaluating heat tolerance within and among species, maximum EHL/MHP becomes particularly informative, with reports of maximum EHL/MHP falling between 1.57 - 2.60 in some passerines (Whitfield et al., 2015), to > 5.0 in a southern African nightjar (O’Connor et al., 2017). One of the major determinants of maximum EHL/MHP appears to be the primary pathway of evaporative heat dissipation: panting, gular flutter or cutaneous evaporation. Species using either gular flutter or cutaneous evaporation as their predominant mechanism for heat dissipation generally exhibit little or no increase in metabolic rate, yet achieve large increases in EWL and thereby attain high EHL/MHP (e.g., Marder and Arieli, 1988; McKechnie et al., 2016; O’Connor et al., 2017). In contrast, species that rely on panting as their primary avenue of EWL typically exhibit larger increases in MHP when actively dissipating heat (Weathers, 1981; Weathers and Greene, 1998). Consequently, when compared at the same T_a_, species that employ gular flutter or cutaneous EWL as their main avenue of heat dissipation tend to have higher EHL/MHP values than species that pant (e.g., Smith et al., 2015; Whitfield et al., 2015; O’Connor et al., 2017). The evolution of evaporative cooling mechanisms minimizing endogenous heat production would have considerable adaptive significance for species subjected to potentially long periods of heat stress, because a reduced internal heat load would require the dissipation of less total heat and, therefore, reduced water demands.

We hypothesized that nocturnal birds have evolved reduced EWL and more efficient evaporative cooling (i.e., higher EHL/MHP at a given T_a_) compared to diurnal species of similar M_b_. We tested three specific predictions regarding EWL and EHL/MHP ratios at T_a_ approaching and exceeding T_b_: 1) nocturnal species display more gradual increases in EWL with increasing T_a_ (i.e., shallower slopes of EWL against T_a_) when the latter exceeds T_b_, 2) nocturnal species have lower EWL at a given T_a_ above normothermic T_b_ and, 3) nocturnal species have higher EHL/MHP at a given T_a_ above normothermic T_b_.

## Materials and methods

### Study species and physiological data

We collated published and unpublished data on T_b_, EWL rates and carbon dioxide production (*V̇CO_2_*) at T_a_ between 10 and 65°C for 39 bird species (Table S1). These data were collected between 2012 and 2015 for an ongoing, collaborative project between multiple laboratories investigating heat tolerance and evaporative cooling efficiency in arid-zone birds spanning three continents. Therefore, all data were collected using similar protocols and under similar experimental conditions (e.g., relatively high flow rates to ensure low humidity in the metabolic chambers; see Whitfield et al., 2015 for details). The inclusion of physiological data from only semi-arid or arid habitats controlled for the potential influence of climate on physiological estimates. Some of the data for several species included here have recently been published, while the remaining data are currently unpublished (Table S1). For the published studies that investigated seasonal differences in heat tolerance, we only used data collected during warm periods (e.g., Noakes et al., 2016; O’Connor et al., 2017). All equipment and experimental protocols used to record T_a_, T_b_, *V̇CO_2_* and EWL rates are described in detail within the recently published studies (Table S1). Additionally, these studies provide details on the thermal equivalents and specific equations used for calculating MHP and EHL.

**Table 1.**
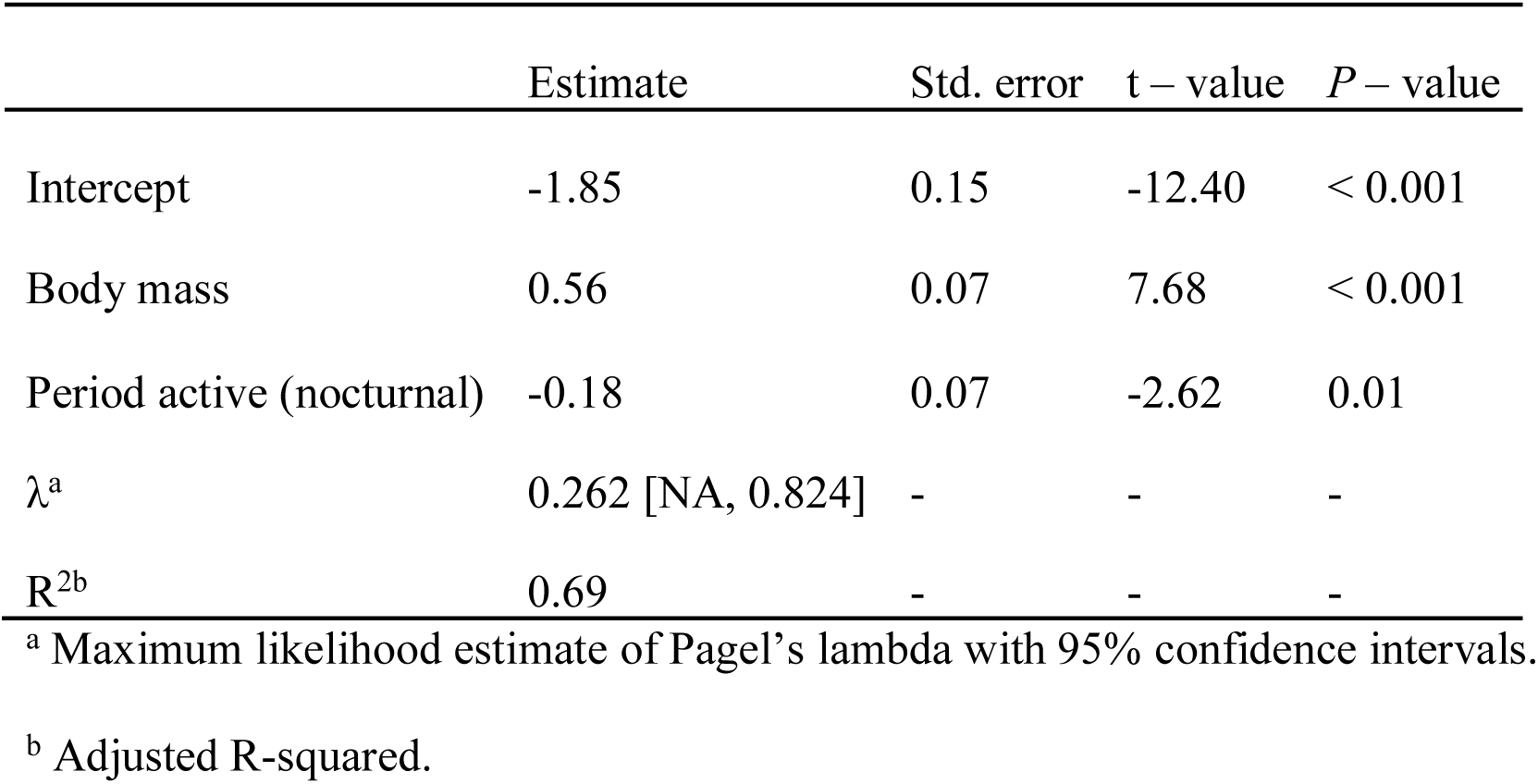
Regression estimates from a phylogenetic generalized least squares model explaining variation in the slopes of evaporative water loss rates at air temperatures between 40 and 46°C in 34 avian species. Predictor variables in the model included log_10_ body mass and period active (diurnal [*N* = 28] *versus* nocturnal [*N* = 6]).

We opted not to include physiological data on nocturnal species from earlier studies because the methodologies and techniques used during open flow-through respirometry can substantially influence physiological estimates (e.g., Hayes et al., 1992; Page et al., 2011; Gerson et al., 2014; Jacobs and McKechnie, 2014). The impact of methodology is clearly demonstrated when considering the relationship between flow rate, humidity and EHL/MHP. Pioneering investigations into heat tolerance of birds used low flow rates when exposing individuals to increasingly high T_a_ (see Lasiewski et al., 1966). Consequently, as birds increased EWL with increasing T_a_, the low flow rates allowed chamber humidity to build up to high levels, impeding birds’ capacity to dissipate heat. Lasiewski et al. (1966) demonstrated that by increasing flow rates, water vapor pressure inside the chamber decreased, allowing birds to tolerate T_a_ values that earlier work had found to be lethal. Previous work by investigators measuring EWL among nocturnal species used low flow rates across all T_a_ (e.g., 830 and 1,262 ml min^-1^ [Ligon, 1968]; 1,500 ml min ^-1^ [Coulombe, 1970]; 2,000 ml min^-1^ [Ganey et al., 1993]), whereas the studies included here all used flow rates ranging from 2,000 up to 30,000 ml min^-1^. Hence, by implementing similar experimental protocols between measurements we were able to minimize noise in our data and permit direct comparisons among species. Two of the nocturnal species included here (*Caprimulgus rufigena* and *Caprimulgus tristigma*) were measured during their diurnal rest phase, whereas the remaining nocturnal species were measured during their nocturnal active phase. However, the influence of the circadian cycle on EWL among birds appears small, with EWL marginally lower during the rest phase, whereas EHL/MHP is seemingly unaffected (Bartholomew and Trost, 1970; Weathers and Caccamise, 1975; Weathers and Schoenbaechler, 1976). Thus, we do not believe that the inclusion of these two caprimulgids significantly influenced our results.

### Data analysis

To quantify variation in EWL and EHL/MHP we fitted the following three models with *period active* representing a two-level categorical predictor (i.e., day or night) and three continuous predictors: *body mass*, the thermal gradient between T_a_ and T_b_ (i.e., *T*_a_ *– T*_b_) and the *body mass:T*_a_ *– T*_b_ interaction term:

(a) Slope of EWL (g hr^-1^ °C^-1^) at T_a_ between 40 and 46°C = body mass + period active,

(b) EWL (g hr^-1^) at 46°C = body mass + period active + T_a_ – T_b_ + body mass:T_a_ – T_b_,

(c) EHL/MHP at 46°C = body mass + period active + T_a_ – T_b_ + body mass:T_a_ – T_b_.

We included *T*_a_ *– T*_b_ as a continuous predictor to control for the possible influence that T_a_ – T_b_ can have on EWL (see Tieleman and Williams, 1999).

All analyses were performed in R v. 3.3.2 (R Core Team 2017). For model (a), we calculated species-specific slopes of EWL at T_a_ between 40 and 46°C (specifically, 39.5°C ≤ T_a_ < 46.5°C) by fitting linear mixed-effect models of EWL rates against T_a_ using the R packages *lme4* (Bates et al., 2015) or *robustlmm* (Koller, 2016). We included bird identity as a random effect to account for repeated measurements within individuals. For models (b) and (c), EWL rates and EHL/MHP ratios represent species averages recorded at T_a_ = 46°C, as this was the best compromise between maximizing sample size and the data available for high T_a_. Given that we included *T*_a_ *– T*_b_ in models (b) and (c), we removed incomplete cases at T_a_ = 46°C when T_b_ was not known due to a bird moving out of range of the antenna receiving the T_b_ signal. Prior to fitting models, we log_10_ transformed EWL slopes at T_a_ between 40 and 46°C, EWL rates at T_a_ = 46°C and *body mass* to correct for skewness in the data. For models (b) and (c), we also mean-centered and standardized the *body mass* and *T*_a_ *– T*_b_ input variables to units of two standard deviations (Gelman, 2008; Schielzeth, 2010). Lastly, we checked for collinearity among our predictors by calculating correlation coefficients (*r*) and variance inflation factors (VIF; maximum *r* = 0.34 and maximum VIF = 2.40).

We analyzed our three models in a phylogenetic generalized least squares (PGLS) regression framework (Symonds and Blomberg, 2014) using the R packages *ape* (Paradis et al., 2004) and *caper* (Orme et al., 2013). We downloaded a subset of 500 phylogenies that included all species in our dataset from http://www.birdtree.org (Jetz et al., 2012) using the Hackett et al. (2008) phylogeny as a back-bone. We subsequently used these 500 phylogenies to obtain a majority consensus tree using “Mesquite” (Madison and Madison, 2017), which was then incorporated into our PGLS models. We followed Revell (2010) and simultaneously estimated the PGLS regression coefficients along with the phylogenetic signal using a maximum likelihood estimation procedure of Pagel’s lambda (λ; Pagel, 1999) bounded by 0 and 1, where 0 indicates phylogenetic independence and 1 indicates that traits follow a Brownian motion model of evolution (Freckleton et al., 2002). The fit of each model was diagnosed by visually inspecting plots of the phylogenetic residuals to assess whether they met the assumptions of normality and homogeneity (Symonds and Blomberg, 2014). Species with a studentized phylogenetic residual > 3.0 or < −3.0 were excluded from the final model as these can overly influence the results of the PGLS regression (Table S1; Jones and Purvis, 1997). Lastly, seven species were represented by one or two individuals at T_a_ = 46°C (Table S1) and we tested for the influence of sample size by running two analyses, one where all species were included and one excluding species for which the data came from fewer than three individuals (McKechnie and Wolf, 2004a; Londoño et al., 2014). The results from these duplicate analyses were quantitatively similar (i.e., estimated effect sizes with standard errors overlapped) and we therefore present the results for the entire data set only.

## Results

Nocturnal species had significantly shallower EWL slopes at T_a_ between 40 and 46°C compared to diurnal species (Table 1). However, among nocturnal birds EWL slopes were noticeably shallower within non-Strigiformes (i.e., caprimulgids and Australian Owlet-nightjars; Figure 1). *Body mass* strongly influenced EWL slopes (Table 1 and Figure 1).

**Fig. 1.**
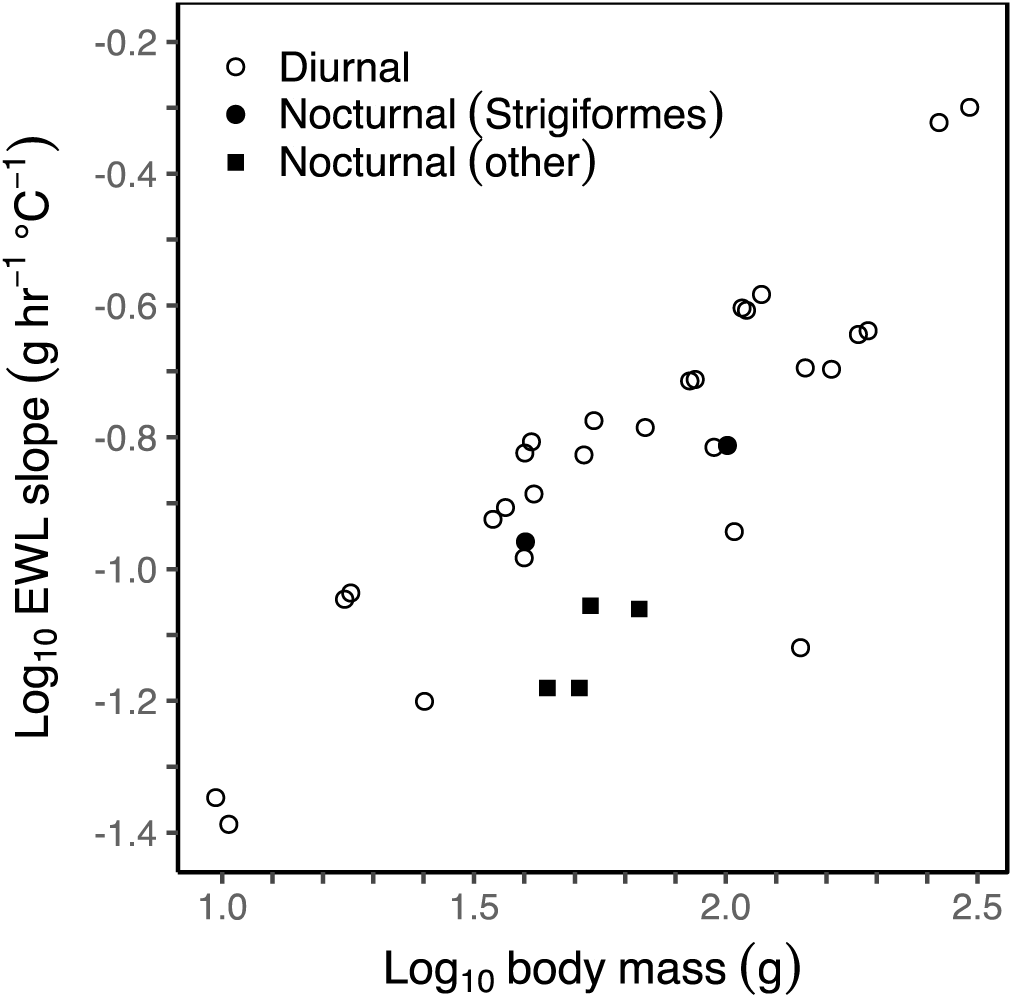
The relationship between the slope of evaporative water loss (EWL) rate at air temperatures between 40 and 46°C with body mass (M_b_) in diurnal (*N* = 28) and nocturnal (*N* = 6) bird species. Nocturnal Strigiformes represent species in the order Strigiformes (i.e., owls) whereas Nocturnal other represent nocturnal species from other orders of nocturnal birds (i.e., nightjars and nighthawks [Caprimulgiformes] and Australian owlet-nightjars [Apodiformes]).

*Body mass* was the most important predictor of EWL at T_a_ = 46°C (Table 2). The regression estimate for nocturnal activity was negative, indicating that nocturnal species had lower EWL than diurnal species of a similar mass (Table 2). However, the effect of nocturnal activity was weak given its relatively small effect size (Table 2). Among nocturnal birds, non-Strigiformes members displayed lower EWL at T_a_ = 46°C compared to diurnal species of comparable M_b_, whereas EWL for Strigiformes (i.e., owls) was generally similar to that of diurnal species of similar size (Figure 2). Consequently, the removal of owls from the data set resulted in nocturnal activity having a stronger effect, although it remained statistically non-significant (*period active nocturnal* = −0.09 ± 0.07, t = −1.28, p = 0.21). Neither *T_a_ – T_b_* nor the interaction between *M_b_* and *T_a_ – T_b_* had a significant influence on EWL at T_a_ = 46°C (Table 2).

**Fig. 2.**
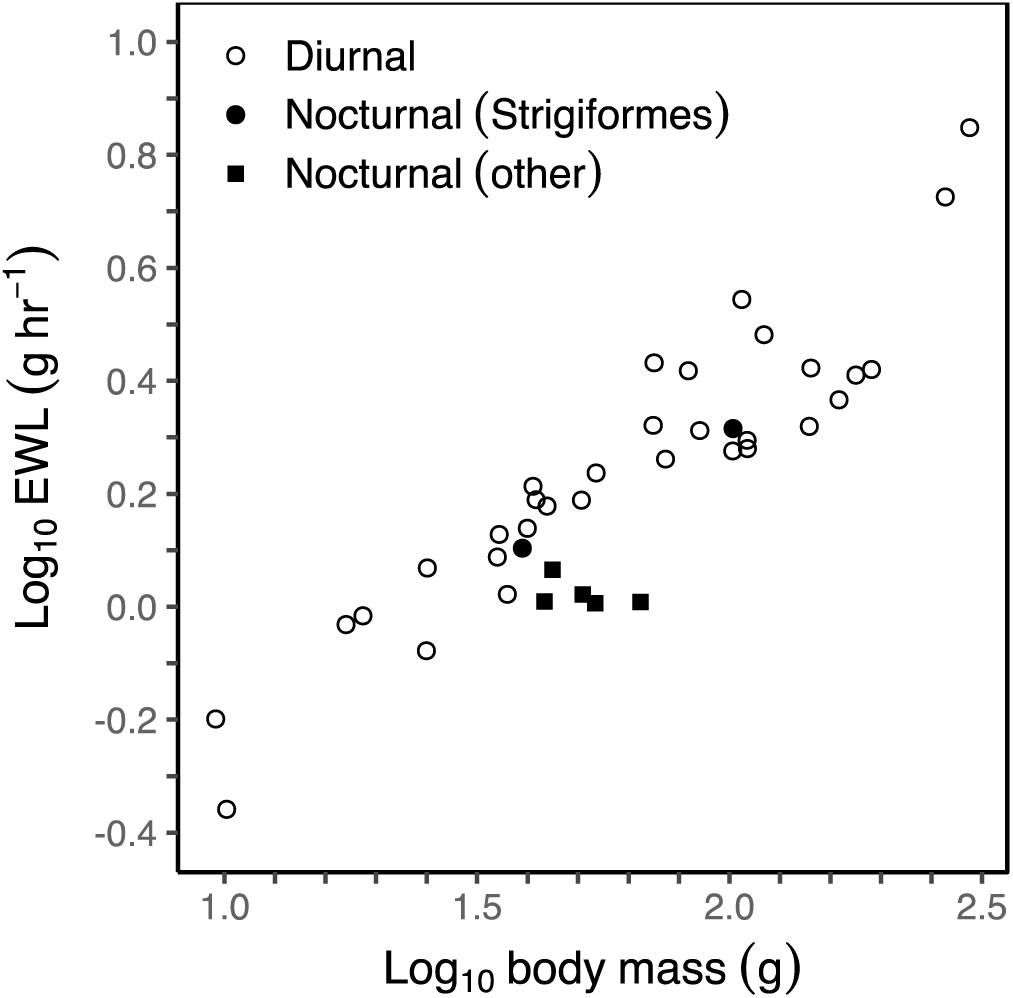
The relationship between evaporative water loss (EWL) rates at an air temperature of 46°C and body mass in diurnal (*N* = 32) and nocturnal (*N* = 7) bird species. Nocturnal Strigiformes represent species in the order Strigiformes (i.e., owls) whereas Nocturnal other represent nocturnal species from other orders of nocturnal birds (i.e., nightjars and nighthawks [Caprimulgiformes] and Australian owlet-nightjars [Apodiformes]).

**Table 2.**
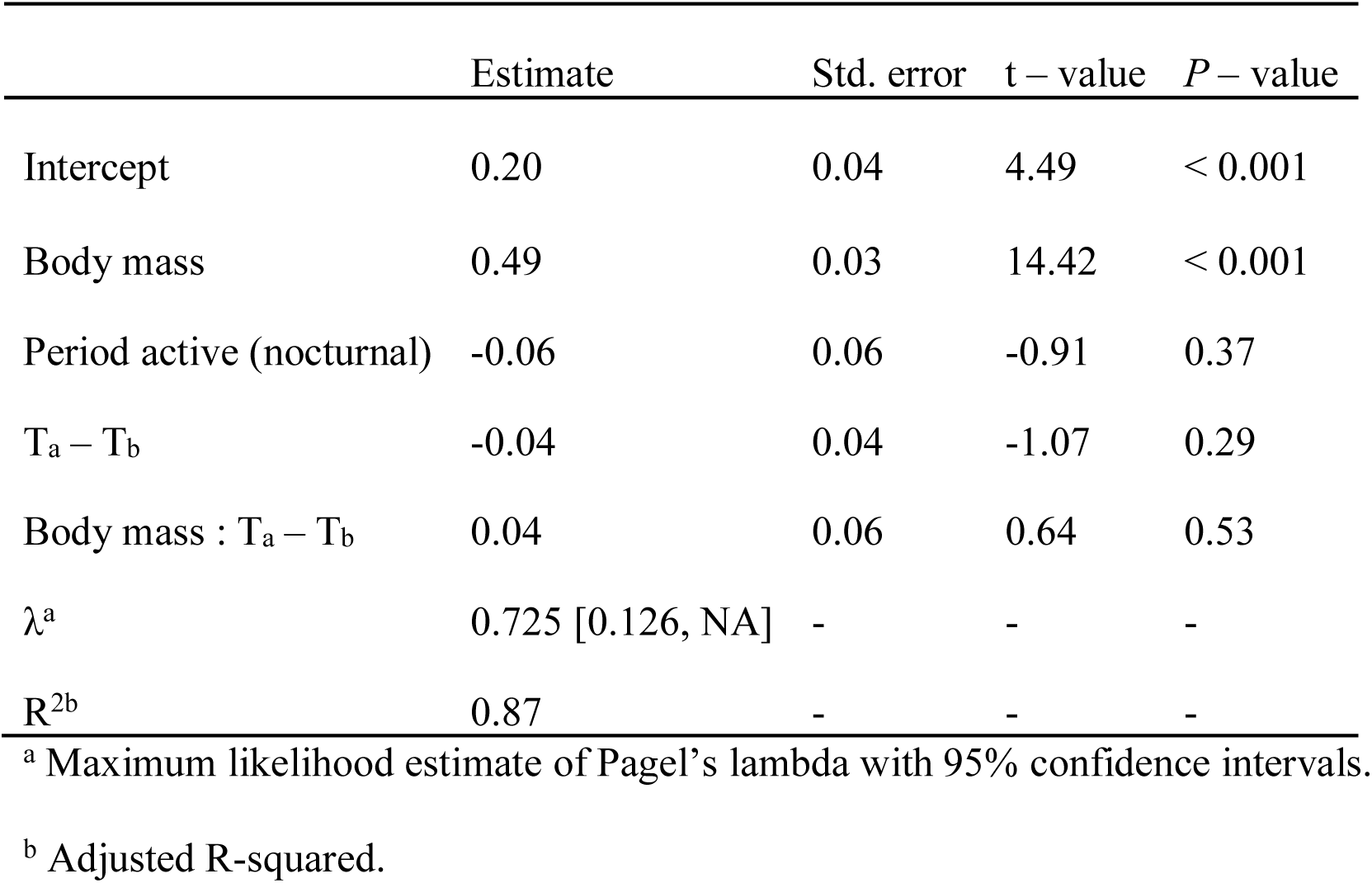
Standardized regression estimates from a phylogenetic generalized least squares model explaining variation in evaporative water loss rates at an air temperature of 46°C for 39 avian species. Predictor variables in the model included log_10_ body mass, period active (diurnal [*N* = 32] *versus* nocturnal [*N* = 7]), the thermal gradient between air temperature and body temperature (T_a_-T_b_) and the interaction between log_10_ body mass and the thermal gradient (body mass:T_a_ – T_b_). Prior to fitting the model, both log_10_ body mass and T_a_ – T_b_ were mean centered and standardized to two standard deviations.

*Body mass* and *T_a_ – T_b_* were the most important predictors of EHL/MHP at T_a_ = 46°C based on the larger standardized effect sizes, with *M_b_* having an overall negative effect and *T_a_ – T_b_* a positive effect (Table 3). Nocturnal activity had a positive effect on EHL/MHP, implying higher EHL/MHP in nocturnal species (Table 3). However, we found considerable variation among nocturnal birds, with owls having lower EHL/MHP ratios compared to non-Strigiformes members (Figure 3). The removal of owls from the data set resulted in *period active* having the largest effect and becoming the most important predictor of EHL/MHP ratios, but again remaining statistically non-significant (*period active nocturnal* = 0.24 ± 0.24, t = 1.03, p = 0.31). Despite many columbids (i.e., doves and pigeons) having comparatively high EHL/MHP ratios at T_a_ = 46°C (Figure 3), all nocturnal birds except Elf Owls maintained a lower T_b_ at T_a_ = 46°C (Figure 4).

**Fig. 3.**
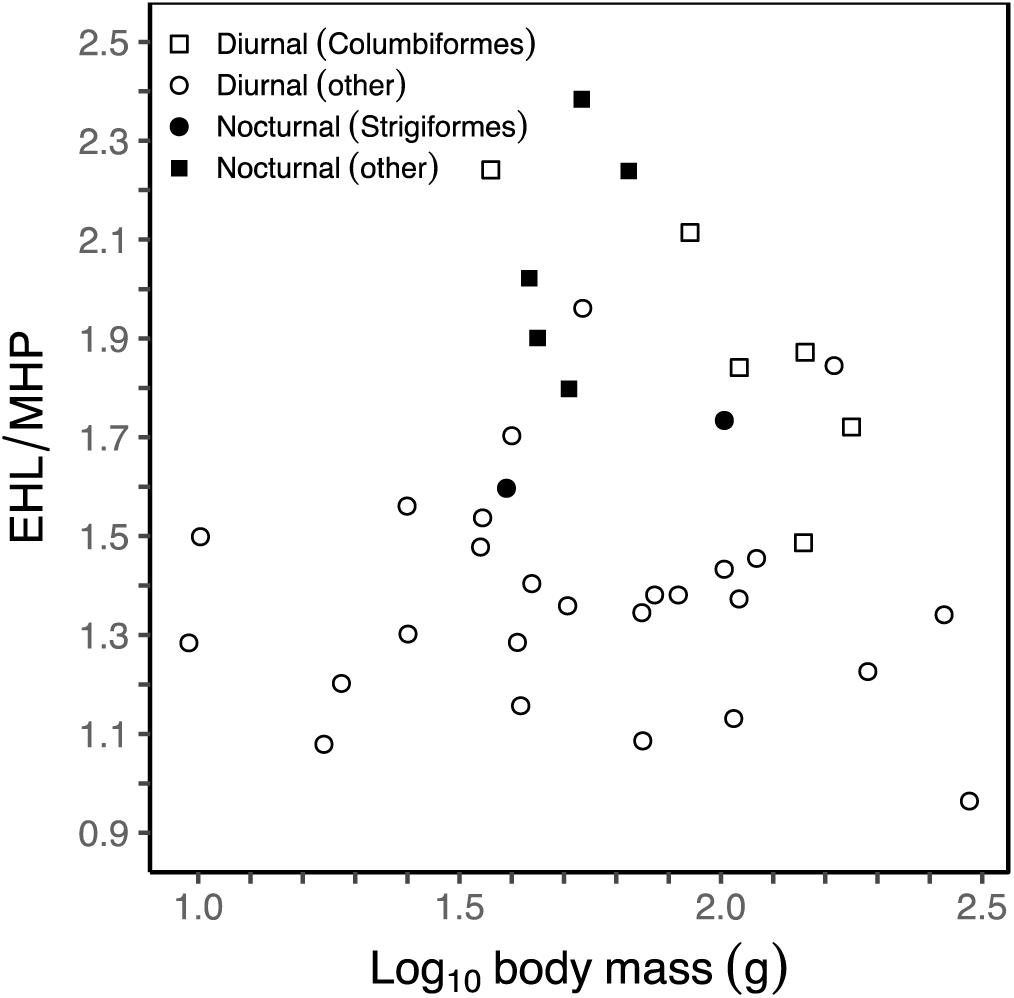
The relationship between the ratio of heat lost through evaporation (EHL) to heat produced through metabolism (MHP; i.e., evaporative cooling efficiency) at an air temperature of 46°C with body mass in diurnal (*N* = 32) and nocturnal (*N* = 7) bird species. Values above one signifies species that dissipated >100% of their metabolic heat through evaporation. Diurnal Columbiformes represent species in the order Columbiformes (i.e., doves and pigeons) whereas Diurnal other represent diurnal species from other orders of diurnal birds. Nocturnal Strigiformes represent species in the order Strigiformes (i.e., owls) whereas Nocturnal other represent nocturnal species from other orders of nocturnal birds (i.e., nightjars and nighthawks [Caprimulgiformes] and Australian owlet-nightjars [Apodiformes]).

**Fig. 4.**
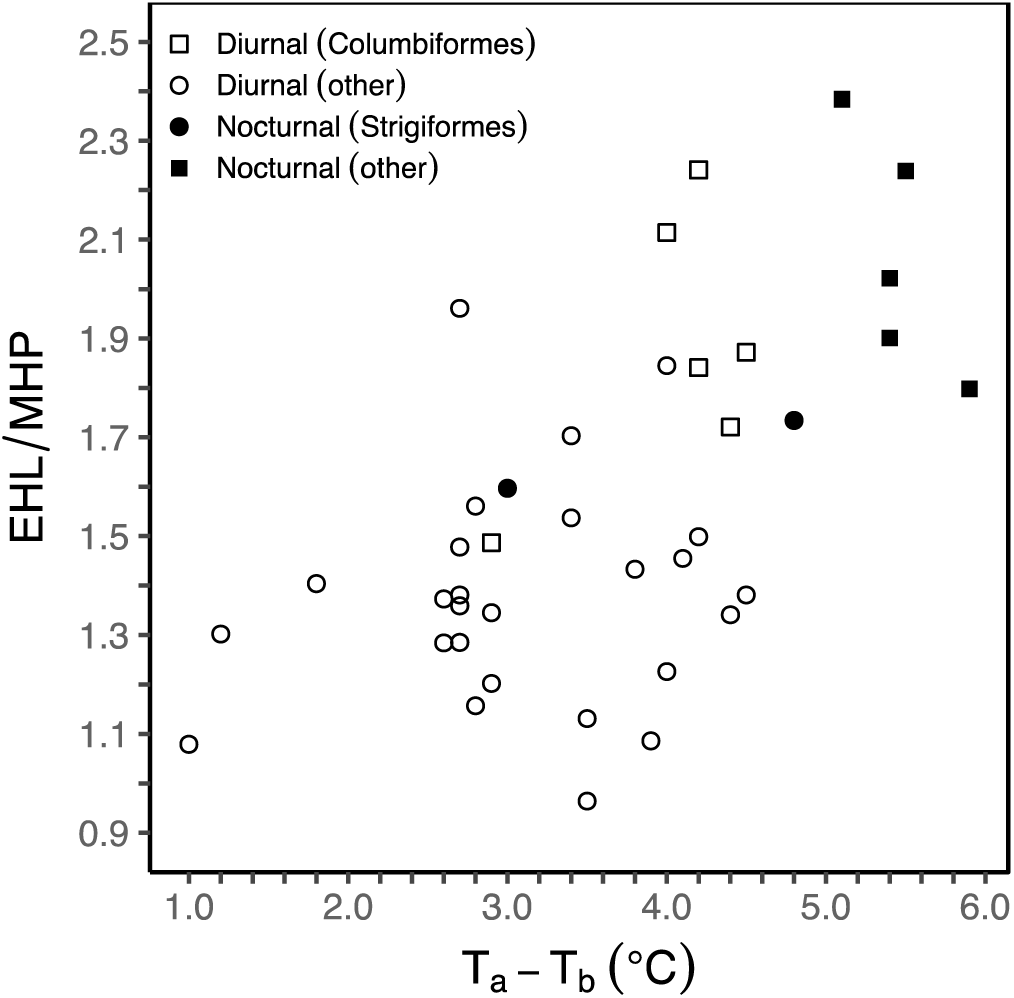
The relationship between the ratio of heat lost through evaporation (EHL) to heat produced through metabolism (MHP; i.e., evaporative cooling efficiency) at an air temperature of 46°C with the thermal gradient between air temperature and body temperature (T_a_ – T_b_) in diurnal (*N* = 32) and nocturnal (*N* = 7) birds. Values above one signifies species that dissipated > 100% of their metabolic heat through evaporation. Diurnal Columbiformes represent species in the order Columbiformes (i.e., doves and pigeons) whereas Diurnal other represent diurnal species from other orders of diurnal birds. Nocturnal Strigiformes represent species in the order Strigiformes (i.e., owls) whereas Nocturnal other represent nocturnal species from other orders of nocturnal birds (i.e., nightjars and nighthawks [Caprimulgiformes] and Australian owlet-nightjars [Apodiformes]).

**Table 3.**
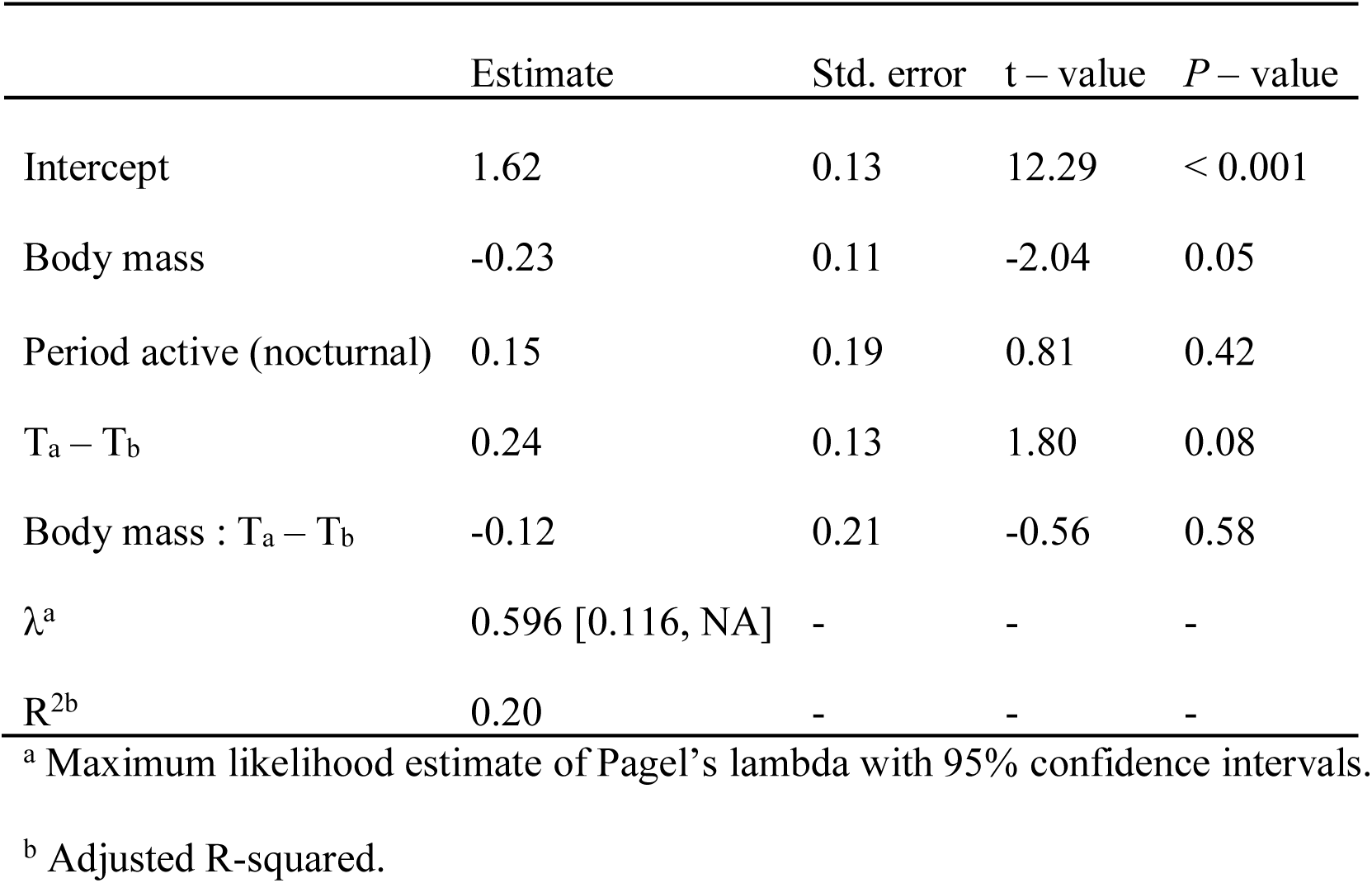
Standardized regression estimates from a phylogenetic generalized least squares model explaining variation in evaporative cooling efficiency at an air temperature of 46°C for 39 avian species. Predictor variables in the model included log_10_ body mass, period active (diurnal [*N* = 32] *versus* nocturnal [*N* = 7]), the thermal gradient between air temperature and body temperature (T_a_ - T_b_) and the interaction between log_10_ body mass and the thermal gradient (body mass:T_a_ – T_b_). Prior to fitting the model, both log_10_ body mass and T_a_ – T_b_ were mean centered and standardized to two standard deviations.

|  | Estimate | Std. error | t – value | P – value |
| --- | --- | --- | --- | --- |
| Intercept | 1.62 | 0.13 | 12.29 | < 0.001 |
| Body mass | -0.23 | 0.11 | -2.04 | 0.05 |
| Period active (nocturnal) | 0.15 | 0.19 | 0.81 | 0.42 |
| $T_a - T_b$ | 0.24 | 0.13 | 1.80 | 0.08 |
| Body mass : $T_a - T_b$ | -0.12 | 0.21 | -0.56 | 0.58 |
| $\lambda^a$ | 0.596 [0.116, NA] | - | - | - |
| $R^{2b}$ | 0.20 | - | - | - |

<sup>a</sup> Maximum likelihood estimate of Pagel's lambda with 95% confidence intervals.
<sup>b</sup> Adjusted R-squared.

## Discussion

Our analysis revealed that increases in EWL with increasing T_a_ are significantly shallower in nocturnal species, but that neither EWL nor EHL/MHP at T_a_ = 46°C are significantly lower. Although our analyses were constrained by the small number of species for which suitable data are currently available, one pattern to emerge concerns differences between the two major radiations of nocturnal birds for which data are available. Whereas the data for caprimulgids and Australian owlet-nightjars appear to be consistent with our predictions of more efficient evaporative cooling in nocturnal birds, the EWL of owls is similar to that of diurnal species. Consequently, the lower *period active* estimates within our EWL models appear to be driven by caprimulgids and Australian Owlet-nightjars, rather than being representative of a generalized evolutionary response to nocturnal activity. Indeed, the removal of owls from the data set resulted in a more negative effect of *period active* on EWL at T_a_ = 46°C, although the effect remained statistically non-significant, likely a corollary of the small sample size.

We did not find consistent support for our prediction that nocturnal species have higher EHL/MHP ratios at a given T_a_ compared to diurnal species of comparable mass. The distinction between nocturnal and diurnal birds was particularly influenced by the high EHL/MHP values of columbids, a result supporting earlier studies that found both caprimulgids and columbids to exhibit high EHL/MHP (e.g., Bartholomew et al., 1962; Dawson and Fisher, 1969; Dawson and Bennett, 1973; Marder and Arieli, 1988; McKechnie et al., 2016a). The extraordinary effectiveness of evaporative cooling among caprimulgids and columbids presumably stems in part from their capacity to dissipate heat by gular flutter and cutaneous evaporation, respectively, which manifests in a rapid elevation in EWL with little or no metabolic cost (Lasiewski and Bartholomew, 1966; Dawson and Fisher, 1969; Marder and Arieli, 1988; Marder et al., 2003; McKechnie and Wolf, 2004b).

Among nocturnal birds, EHL/MHP was lower in owls compared to caprimulgids and Australian Owlet-nightjars. Indeed, when owls were excluded from the data set, both the effect size and relative importance for *period active* increased, suggesting that the inclusion of data for owls reduced the overall effect. These observed differences may represent the varying contributions of panting to gular flutter among these nocturnal groups. When exposed to increasing heat loads under laboratory conditions, panting and gular flutter rates became synchronized with each other in Barn Owls (*Tyto alba*), Burrowing Owls (*Athene cunicularia*) and Great Horned Owls (*Bubo virginianus*; Bartholomew et al., 1968; Coulombe, 1970). The presence of rapid panting, even if concurrent with the metabolically efficient mechanism of gular fluttering, should increase a bird’s total heat load because of the consequential increase in endogenous heat production associated with moving the thoraco-abdominal structure (Dawson and Whittow, 2000). In contrast, gular flutter appears to be asynchronous and occurs at a greater frequency than breathing rate among caprimulgids (Lasiewski and Bartholomew, 1966; Bartholomew et al., 1968; Dawson, 1982; Dawson and Whittow, 2000). Rapid vibrations of the gular region are believed to have a lower energetic cost than panting because the anatomical structures involved are moved more efficiently than those used for panting (Lasiewski and Bartholomew, 1966). Additionally, the enlarged, highly vascularized gular area of caprimulgids provides a larger surface area for evaporative heat dissipation, further enhancing evaporative cooling efficiency (Lasiewski and Bartholomew, 1966).

We acknowledge that our sample size of 39 species is relatively small for a phylogenetic analysis, particularly when compared to past avian EWL reviews (e.g., Crawford and Lasiewski, 1968, n = 53; Williams, 1996, n = 102). Furthermore, our sample of nocturnal species only represented 17.9% of the total data at T_a_ = 46°C. However, our data set addresses an important, long standing issue among biological and ecological studies, that of biological *versus* statistical significance (Yoccoz, 1991; Johnson, 1999; Martínez-Abraín, 2008). Statistical significance is strongly dependent on sample size and frequently investigators obtain a result that is biologically important but statistically non-significant, or *vice versa* (Gerrodette, 2011). Currently, the small and unbalanced sample size of our nocturnal species (two Strigiformes and five non-Strigiformes) precludes rigorous statistical comparisons between these two nocturnal evolutionary radiations, but the observed variation in EWL and EHL/MHP between them is arguably biologically relevant with regards to water conservation and dehydration avoidance. However, we argue that the inclusion of data for nocturnal birds from the literature, where birds were exposed to higher humidity due to lower flow-rates, would result in greater noise in our dataset. Indeed, earlier EWL reviews that collated data across myriad studies acknowledged the possible addition of extraneous variation due to differing experimental techniques and conditions (Crawford and Lasiewski, 1968; Williams, 1996; Tieleman and Williams, 1999).

The apparent disparity in EWL and EHL/MHP among our nocturnal birds likely reflects the ecological differences between these groups. During their diurnal rest-phase, owls often select cavities and/or shaded sites with cooler microclimates (Barrows, 1981; Hardy and Morrison, 2001; Charter et al., 2010), a behavior presumably necessary because of a physiological constraint in their capacity to reduce EWL at high T_a_. Ganey (2004), for example, suggested that Spotted Owls (*Strix occidentalis*) select nest sites with thermal properties conferring a positive water balance. Elf Owls nesting in the Sonoran Desert of Arizona, USA, prefer north-facing cavities inside saguaro cacti where maximum internal cavity temperatures may be 2.8 - 6.0°C below maximum T_a_ (Soule, 1964; Hardy and Morrison, 2001), a thermal gradient that would reduce total heat loads and, consequently, EWL. However, it is noteworthy that temperatures inside refugia still do, on occasion, approach or exceed T_b_ (Soule, 1964; Ligon, 1968). Interestingly, Australian Owlet-nightjars also make use of cavities during the diurnal period (Doucette et al., 2011), but their physiological responses to heat appear more similar to that of caprimulgids and may reflect these two groups sharing a more recent common ancestor (Hackett et al., 2008). The occupancy of unshaded sites with high operative temperatures by some caprimulgids is no doubt possible through their ability to minimize EWL while maintaining high EHL/MHP ratios via a metabolically efficient mechanism of heat dissipation, namely gular flutter. Furthermore, the use of open nest sites by ground nesting species is believed to aid in predator detection and increased survival (Amat and Masero, 2004). Thus, the ability of caprimulgids to roost and nest in the open may translate into increased fitness.

## Conclusions

We found limited evidence for more economical evaporative cooling in nocturnal species. Although slopes of EWL were significantly shallower in nocturnal birds, EWL and EHL/MHP at a given T_a_ were not. Our data do, however, suggest considerable variation in physiological responses to high T_a_ between owls and caprimulgids. When compared to diurnal species of similar mass, caprimulgids and Australian Owlet-nightjars had lower EWL rates, whereas owls displayed EWL rates comparable to diurnal species. Despite having higher EWL rates, owls tended to have lower EHL/MHP ratios compared to caprimulgids and Australian Owlet-nightjars. This trend may represent an increased contribution of panting relative to gular fluttering among owls, resulting in increased MHP relative to EHL. Admittedly based on a small sample size, our results allude to a biologically important difference among nocturnal birds in their capacity to conserve water at high T_a_. Barrows (1981), for instance, observed Spotted Owls in the wild exhibiting signs of heat stress at moderate T_a_ of 29 to 31°C, despite these birds occupying roost sites that were 1 – 6°C cooler than the surrounding open areas. Given that heat waves are predicted to increase in occurrence, duration and intensity throughout the 21^st^ century (Meehl and Tebaldi, 2004), current refugia used by owls that provide favorable microclimates may no longer adequately buffer them from heat stress because of more frequent and longer lasting periods when T_a_ exceeds heat stress inducing temperatures. Owls, therefore, may become more susceptible to dehydration risk during extreme weather events compared to other nocturnal birds. More data for nocturnal species could provide additional support as to whether owls indeed lack physiological adaptations for reducing EWL. Similarly, investigations on physiological adaptations conferring water conservation among mammals would be worthwhile given their use of various shelter types (e.g., underground burrows, tree cavities or leaf nests above ground; Reichman and Smit, 1990).

## List of symbols and abbreviations

EHL: evaporative heat loss
EWL: evaporative water loss
M_b_: body mass
MHP: metabolic heat production
T_a_: air temperature
T_b_: body temperature
PGLS: phylogenetic generalized least squares
*V̇CO*_2_: carbon dioxide production

## Acknowledgements

This study represents the culmination of several years of data collection and we are grateful for all the assistants that helped with field work. Michelle Thompson assisted with ggplot2 code for figure construction.

## Author contributions

B.O.W. and A.E.M. designed the study. R.S.O, B.S., W.A.T., A.R.G., B.O.W. and A.E.M. collected data. R.S.O and B.S. analyzed the data. R.S.O. and A.E.M. wrote the original draft. B.S., W.A.T., A.R.G., R.M.B and B.O.W reviewed and edited the manuscript.

