## Supplementary Materials for "Avian thermoregulation in the heat: is evaporative cooling more economical in nocturnal birds?"

**Table S1.** Avian species and data used for phylogenetically-informed analyses on the slope of evaporative water loss rates at air temperatures ($T_a$) between 40 and 46°C (EWL slope), evaporative water loss rates at 46°C (EWL) and the ratio of evaporative heat loss and metabolic heat production at 46°C (EHL/MHP). $T_a$-$T_b$ represents the thermal gradient between $T_a$ and body temperature ($T_b$) at 46°C. $M_b$ slope represents the average body mass for a species included in the EWL slope analysis and $M_b$ EWL represents the average body mass for a species included in the EWL and EHL/MHP analyses. Source represents the published articles from which data were derived. Values in brackets are the number of individuals included in an analysis for a species (e.g., 26 Abert's Towhees were included in the EWL slope analysis and 9 in the EWL and EHL/MHP analyses).

| Species | Period[a] active | $M_b$ (g) slope | EWL slope (g $hr^{-1}$ $°C^{-1}$) | $M_b$ (g) EWL | EWL (g $hr^{-1}$) | EHL/MHP | $T_a$-$T_b$ (°C) | Source |
|---|---|---|---|---|---|---|---|---|
| Abert's Towhee *Melozone aberti* | D | 41.6 | 0.130 [26] | 41.4 | 1.547 [9] | 1.157 | 2.8 | Smith et al. 2017 |
| African Cuckoo *Cuculus gularis* | D | 109.9 | 0.247 [6] | 108.4 | 1.971 [3] | 1.373 | 2.6 | Smit et al. 2018 |
| Apostlebird *Struthidea cinerea* | D | 117.7 | 0.261 [7] | 117.0 | 3.032 [3] | 1.455 | 4.1 | McKechnie et al. 2017 |
| Australian Owlet-nightjar *Aegotheles cristatus* | N | 44.2 | 0.066 [22] | 44.6 | 1.162 [8] | 1.901 | 5.4 | Talbot et al. 2017 |
| Burchell's Sandgrouse *Pterocles burchelli* | D | 191.7 | 0.230 [16] | 191.2 | 2.631 [5] | 1.226 | 4.0 | McKechnie et al. 2016 |
| Burchell's Starling *Lamprotornis australis* | D | 107.7 | 0.249 [7] | 105.8 | 3.499 [3] | 1.131 | 3.5 | Smit et al. 2018 |
| Cactus Wren *Campylorhynchus brunneicapillus* | D | 34.5 | 0.119 [21] | 34.7 | 1.225 [9] | 1.478 | 2.7 | Smith et al. 2017 |

| Species | Period[a] active | $M_b$ (g) slope | EWL slope (g $hr^{-1}$ °$C^{-1}$) | $M_b$ (g) EWL | EWL (g $hr^{-1}$) | EHL/MHP | $T_a$-$T_b$ (°C) | Source |
|---|---|---|---|---|---|---|---|---|
| Cape Glossy Starling *Lamprotornis nitens* | D | NA | NA | 70.9 | 2.703 [1] | 1.086 | 3.9 | *Unpublished data* |
| Chestnut-crowned Babbler *Pomatostomus ruficeps* | D | 52.3 | 0.149 [17] | 50.9 | 1.544 [8] | 1.359 | 2.7 | McKechnie et al. 2017 |
| Common Poorwill[b] *Phalaenoptilus nuttallii* | N | 43.7 | 0.116 [26] | 43.0 | 1.022 [5] | 2.022 | 5.4 | Talbot et al. 2017 |
| Crested Pigeon *Ocyphaps lophotes* | D | 183.5 | 0.227 [36] | 177.9 | 2.571 [9] | 1.721 | 4.4 | McKechnie et al. 2016b |
| Curve-billed Thrasher *Toxostoma curvirostre* | D | 69.2 | 0.164 [16] | 70.6 | 2.095 [4] | 1.345 | 2.9 | Smith et al. 2017 |
| Elf Owl *Micrathene whitneyi* | N | 40.0 | 0.110 [12] | 38.9 | 1.270 [3] | 1.597 | 3.0 | *Unpublished data* |
| Freckled Nightjar *Caprimulgus tristigma* | N | 67.3 | 0.087 [16] | 66.6 | 1.019 [13] | 2.239 | 5.5 | O'Connor et al. 2017 |
| Galah *Eolophus roseicapilla* | D | 265.3 | 0.476 [7] | 267.5 | 5.316 [3] | 1.341 | 4.4 | McWhorter et al. 2018 |
| Gambel's Quail *Callipepla gambelii* | D | 162.1 | 0.201 [17] | 164.8 | 2.325 [6] | 1.845 | 4.0 | Smith et al. 2015 |
| Grey Butcherbird *Cracticus torquatus* | D | 84.8 | 0.193 [15] | 82.9 | 2.617 [4] | 1.381 | 2.7 | McKechnie et al. 2017 |

| Species | Period[a] active | $M_b$ (g) slope | EWL slope (g $hr^{-1}$ °$C^{-1}$) | $M_b$ (g) EWL | EWL (g $hr^{-1}$) | EHL/MHP | $T_a$-$T_b$ (°C) | Source |
|---|---|---|---|---|---|---|---|---|
| House Finch *Haemorhous mexicanus* | D | 18.0 | 0.092 [32] | 18.8 | 0.964 [8] | 1.202 | 2.9 | Smith et al. 2017 |
| Laughing Dove *Spilopelia senegalensis* | D | 87.0 | 0.194 [17] | 87.3 | 2.051 [4] | 2.114 | 4.0 | McKechnie et al. 2016b |
| Lesser Goldfinch *Spinus psaltria* | D | 9.7 | 0.045 [18] | 9.6 | 0.633 [4] | 1.284 | 2.6 | Smith et al. 2017 |
| Lesser Nighthawk *Chordeiles acutipennis* | N | 51.2 | 0.066 [15] | 51.2 | 1.051 [3] | 1.798 | 5.9 | Talbot et al. 2017 |
| Lilac-breasted Roller *Coracias caudatus* | D | 94.8 | 0.153 [7] | 101.5 | 1.887 [4] | 1.433 | 3.8 | Smit et al. 2018 |
| Marico Flycatcher *Melaenornis mariquensis* | D | NA | NA | 25.2 | 1.171 [1] | 1.302 | 1.2 | *Unpublished data* |
| Mourning Dove *Zenaida macroura* | D | 104.0 | 0.114 [41] | 108.5 | 1.906 [8] | 1.841 | 4.2 | Smith et al. 2015 |
| Mulga Parrot *Psephotellus varius* | D | 54.7 | 0.168 [16] | 54.4 | 1.725 [7] | 1.961 | 2.7 | McWhorter et al. 2018 |
| Namaqua Dove *Oena capensis* | D | 36.5 | 0.124 [10] | 36.3 | 1.052 [5] | 2.241 | 4.2 | McKechnie et al. 2016b |
| Northern Cardinal *Cardinalis cardinalis* | D | 39.9 | 0.150 [10] | 40.8 | 1.634 [2] | 1.285 | 2.7 | Smith et al. 2017 |

| Species | Period[a] active | $M_b$ (g) slope | EWL slope (g $hr^{-1}$ °$C^{-1}$) | $M_b$ (g) EWL | EWL (g $hr^{-1}$) | EHL/MHP | $T_a$-$T_b$ (°C) | Source |
|---|---|---|---|---|---|---|---|---|
| Pyrrhuloxia *Cardinalis sinuatus* | D | NA | NA | 35.0 | 1.343 [1] | 1.537 | 3.4 | Smith et al. 2017 |
| Red-billed Buffalo-Weaver[b] *Bubalornis niger* | D | 71.1 | 0.030 [5] | 74.7 | 1.826 [1] | 1.381 | 4.5 | *Unpublished data* |
| Red-billed Spurfowl *Pternistis adspersus* | D | 305.0 | 0.502 [4] | 298.9 | 7.050 [2] | 0.964 | 3.5 | *Unpublished data* |
| Ring-necked Dove *Streptopelia capicola* | D | 140.7 | 0.076 [15] | 143.9 | 2.086 [4] | 1.487 | 2.9 | McKechnie et al. 2016b |
| Rufous-cheeked Nightjar *Caprimulgus rufigena* | N | 53.9 | 0.088 [16] | 54.2 | 1.014 [8] | 2.384 | 5.1 | O'Connor et al. 2017 |
| Scaly-feathered Weaver *Sporopipes squamifrons* | D | 10.3 | 0.041 [14] | 10.1 | 0.438 [6] | 1.499 | 4.2 | Whitfield et al. 2015 |
| Sociable Weaver *Philetairus socius* | D | 25.2 | 0.063 [14] | 25.1 | 0.836 [5] | 1.561 | 2.8 | Whitfield et al. 2015 |
| Spiny-cheeked Honeyeater *Acanthagenys rufogularis* | D | 41.1 | 0.156 [16] | 43.5 | 1.507 [1] | 1.404 | 1.8 | McKechnie et al. 2017 |
| Western Screech Owl *Megascops kennicottii* | N | 100.7 | 0.154 [13] | 101.6 | 2.068 [7] | 1.734 | 4.8 | *Unpublished data* |
| White-browed Sparrow-Weaver *Plocepasser mahali* | D | 39.8 | 0.104 [10] | 39.8 | 1.377 [10] | 1.703 | 3.4 | Noakes et al. 2016 |

| Species | Period[a] active | $M_b$ (g) slope | EWL slope (g $hr^{-1}$ °$C^{-1}$) | $M_b$ (g) EWL | EWL (g $hr^{-1}$) | EHL/MHP | $T_a$-$T_b$ (°C) | Source |
|---|---|---|---|---|---|---|---|---|
| White-winged Dove *Zenaida asiatica* | D | 143.8 | 0.202 [34] | 145.0 | 2.646 [7] | 1.872 | 4.5 | Smith et al. 2015 |
| Yellow-plumed Honeyeater *Ptilotula ornata* | D | 17.5 | 0.090 [11] | 17.4 | 0.930 [3] | 1.079 | 1.0 | McKechnie et al. 2017 |

[a] D = Diurnal, N = Nocturnal

[b] The Common Poorwill and Red-billed Buffalo-weaver were removed from the final EWL slope regression model because these species had studentized phylogentic residuals > 3 or < -3.
